# DeepImageTranslator: a free, user-friendly graphical interface for image translation using deep-learning and its applications in 3D CT image analysis

**DOI:** 10.1101/2021.05.15.444315

**Authors:** Run Zhou Ye, Christophe Noll, Gabriel Richard, Martin Lepage, Éric E. Turcotte, André C. Carpentier

## Abstract

**Objectives:** The advent of deep-learning has set new standards in an array of image translation applications. At present, the use of these methods often requires computer programming experience. Non-commercial programs with graphical interface usually do not allow users to fully customize their deep-learning pipeline. Therefore, our primary objective is to provide a simple graphical interface that allows students and researchers with no programming experience to easily create, train, and evaluate custom deep-learning models for image translation. We also aimed to test the applicability of our tool (the DeepImageTranslator) in two different tasks: semantic segmentation and noise reduction of CT images.

**Methods:** The DeepImageTranslator was implemented using the Tkinter library; backend computations were implemented using Pillow, Numpy, OpenCV, Augmentor, Tensorflow, and Keras libraries. Convolutional neural networks (CNNs) were trained using DeepImageTranslator and assessed with three-way cross-validation. The effects of data augmentation, deep-supervision, and sample size on model accuracy were also systematically assessed.

**Results:** The DeepImageTranslator a simple tool that allows users to customize all aspects of their deep-learning pipeline, including the CNN, the training optimizer, the loss function, and the types of training image augmentation scheme. We showed that DeepImageTranslator can be used to achieve state-of-the-art accuracy and generalizability in semantic segmentation and noise reduction. Highly accurate 3D segmentation models for body composition can be obtained using training sample sizes as small as 17 images. Therefore, for studies with small datasets, researchers can randomly select a very small subset of images for manual labeling, which can then be used to train a specialized CNN model with DeepImageTranslator to fully automate segmentation of the entire dataset, thereby saving tremendous time and effort.

**Conclusions:** An open-source deep-learning tool for accurate image translation with a user-friendly graphical interface was presented and evaluated. This standalone software can be downloaded for Windows 10 at: https://sourceforge.net/projects/deepimagetranslator/

## INTRODUCTION

Image translation or transformation is an important and challenging task in many areas of clinical and fundamental sciences. Since the introduction of convolutional neural networks (CNN), generations of CNN architectures have been designed and have achieved state-of-the-art performance in image translation tasks, such as semantic segmentation [1–8], noise reduction [9–11], and image synthesis [12, 13].

Nevertheless, the application of deep-learning methods for image translation can be difficult for scientists with no computer programming experience. In general, deep-learning pipelines are created using custom-implemented codes. Existing software programs for deep-learning-based image analysis that allow users to build, train, and evaluate custom CNNs, such as Niftynet [14], are mostly accessed through a command-line interface. On the contrary, non-commercial open-source programs that interacts with users with a graphical interface, such as the ImageJ implementation of U-net [15] or ilastik [16], do not allow users to customize their CNN (*e.g.* adjusting the number of channel/layers, specifying input image resolution), the training optimizer, the loss function, and the use of different training image augmentation scheme. Therefore, our primary objective is to create a user-friendly graphical interface that allow students and researchers to easily implement, train, and test custom deep-learning pipelines for image translation. Our secondary objective is to verify the applicability of our tool in two different image translation tasks using CT images.

One specific use of the CNNs is in semantic segmentation of CT images for assessment of body composition. For example, CT segmentation is critical for precise quantification of different adipose tissue compartments to provide useful information for fundamental research in metabolic syndrome. However, most of existing implementations of deep-learning methods in abdominal CT segmentation were made for clinical research using 2D single-slice images from cohorts of thousands of patients, which is not applicable in small scale studies. Furthermore, studies of 3D volumetric segmentation for body composition are also scarce. Therefore, we also aimed to evaluate the practical application of the DeepImageTranslator in our small dataset of 524 volumetric CT images from 5 subjects, while also assessing the effects of data augmentation, deep-supervision, and sample size on model accuracy.

Here, we present the various features of DeepImageTranslator and used CT images to evaluate the performance of our software in two image translation tasks, namely semantic image segmentation and noise reduction. This open-source tool is freely available to the general scientific community and can be installed on a Windows computer and used without any prior knowledge of computer programming.

## METHODS

### DeepImageTranslator software development

DeepImageTranslator is a user-friendly tool freely available at: https://sourceforge.net/projects/deepimagetranslator/ for Windows. It is written in Python 3.8 and distributed under the GNU General Public License (version 3.0). Its graphical user interface was developed using the Tkinter library and allows one to easily load images into the software, customize hyperparameters, and save trained models and predicted target images on the user’s hard drive. Image generation and augmentation prior to training were achieved using codes implemented with Pillow, Numpy, OpenCV, and Augmentor libraries. Model training and image prediction were implemented using Tensorflow and Keras.

### CT image acquisition

We evaluated our pipeline using data from 5 subjects who underwent PET/CT examinations as part of a different study in our laboratory, which will be published elsewhere. Whole body CT scans (16 mAs) were performed with a 16-slice PET/CT scanner (Philips Gemini GXL; Philips, Eindhoven, The Netherlands); image reconstruction was done with a row action maximum likelihood algorithm without sinogram rebinning. Final scans had a diameter of 60 cm, axial field of view of 18 cm, and isotropic voxel size of 4 mm. From each subject, approximately 50 consecutive axial images were selected from the level of the sacroiliac joint to the level of the portal hepatic vein bifurcation, which all contained a visceral adipose tissue compartment. From the 5 study participants (263 images), we selected the most obese (53 images) and the leanest individual (42 images) as our test subjects, leaving the other 3 subjects (165 images) for model training and validation. For the protocol involving human subjects, which received approval from the Human Ethics Committee of Centre de recherche du CHUS, informed written consent was obtained from all participants in accordance with the Declaration of Helsinki.

### Manual CT image segmentation

Manual segmentation of CT images was done using a semi-automatic approach with the GIMP software (GNU Image Manipulation Program). Segmentation maps were independently examined and validated by one of the co-authors. Briefly, the subcutaneous adipose tissue (ScAT) compartment was first delineated and colored in red (RGB (red, green, blue) = (255,0,0)). Then, the visceral adipose tissue (VAT) compartment was singled out (using pixel intensity thresholding) and colored in green (RGB = (0,255,0)). The rest of the tissues (including muscles, bones, and intraabdominal organs) were labelled in blue (RGB = (0,0,255)), while the background was labelled in black (RGB = (0,0,0)). Thus, segmentation maps for each of the three compartments were stored in distinct channels of the RGB color space.

### Model training and assessment

Convolutional neural networks were constructed with the DeepImageTranslator pipeline. For model hyperparameters, we used 5 U-net layers, 16 channels following the first convolution, and a batch number of 1.

Pixel values of the input CT images and of their corresponding segmentation maps were automatically normalized to values between 0 and 1 by the DeepImageTranslator before feeding them to the model for training. Therefore, the semantic segmentation task becomes a simple classification task whereby for each pixel in the input image, the model must determine whether the underlining region belongs to the subcutaneous adipose tissue, visceral adipose tissue, or lean tissue compartment by assigning a value of 1 to the red, green, or blue channel, respectively and, alternatively, a value of 0 to all channels if the pixel belongs to the background.

We employed the Adaptive Moment Estimation (Adam) optimizer with a learning rate set at 0.001. Binary cross-entropy loss was computed and used as the loss function. For assessment of model performance, we used the Dice coefficient, sensitivity, specificity, and pixel area difference. For a given channel *c*, the Dice coefficient (DC) between the true target (*T*) and the model predicted target (*P*) was calculated for a given volumetric scan using the following formula:

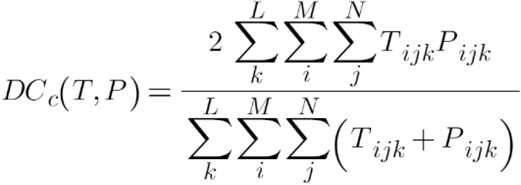

*T_ijk_* and *P_ijk_* represent the value of the pixel located on row *i* and column *j* of slice *k* of the true target and the model predicted target, respectively, which is, after normalization, wither 0 or 1. *M* and *N* are the height and width of the segmentation map, in pixels; *L* represents the total number of slices. Similarly, the sensitivity (*Sn*), specificity (*Sp*), and area difference (*AD*) were determined using the following three equations, respectively:

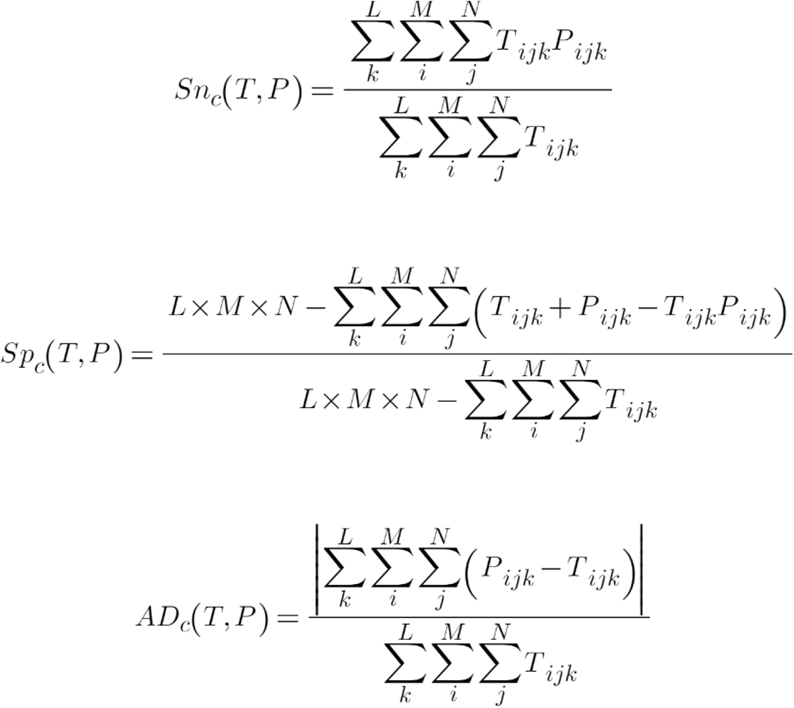

The structural similarity index measure (SSIM) [17], the peak signal-to-noise ratio (PSNR), mean average error (MAE) and mean squared error (MSE) were calculated using Tensorflow 2.4.0.

### Data augmentation

Training image and target augmentation was performed using random perspective scaling, rotation, translation, and flipping. Perspective scaling was accomplished by randomly choosing new image vertices in a square area with side length of 60 pixels centered at the vertices of the original image. The new image formed by these new vertices was then scaled back to the dimension of the original image. Rotation was done by randomly rotating the original image by a maximum of 70 degrees; translation was performed by random translation along the x- and the y-axes by a maximum of 30 pixels; finally, images were randomly flipped along the x- and the y-axes.

Gaussian noise was introduced by randomly adding noise signals to pixels of the original image. The intensity of the noise signal was randomly selected and follows a standard normal distribution (*i.e.* Gaussian distribution centered at 0 with a standard deviation of 1). For a given image, the probability of applying Gaussian noise to a given pixel was randomly chosen between 0 and a user-determined probability threshold. This threshold was set to 30% for our image denoising training experiments. For some of the experiments designed to test model generalizability (see Results), we increased this threshold to 50% (a 60% increase).

### Statistical analyses

Statistical analyses were carried out using GraphPad Prism version 9. Three-way and two-way ANOVAs were carried out to determine the effects of various techniques on model accuracy. P-values smaller than 0.05 were considered significant and were corrected for multiple testing when applicable.

### Data availability

The source code for the DeepImageTranslator is publicly available at: https://github.com/runzhouye/DeepImageTranslator

The compiled standalone software is available for Window10 at: https://sourceforge.net/projects/deepimagetranslator/

The datasets generated during and/or analyzed during the current study are available at: https://figshare.com/s/b9caed5d78f9debdcd21

## RESULTS

### DeepImageTranslator

All the features of DeepImageTranslator can be accessed through its graphical user interface (Fig.1), which includes: a main window for the visualization of input, target, and network-predicted images in the training, validation, and test/translation sets (Fig.1a); a window for selecting the type of model optimizer (*i.e.* Adam, RMSprop, SDG, Adagrad, and Adadelta), loss function (*i.e.* binary cross-entropy, categorical cross-entropy, MSE, MAE, and sparse categorical cross-entropy), and training metrics, as well as the number of training epochs and batch size (Fig.1b); a window for neural network construction, which allows full customization of input and target dimensions, number of layers, number of channels at the end of the first convolution, and the inclusion of a deep-supervision layer (Fig.1c); a window to monitor the training process (Fig.1d). A window for the selection of data augmentation schemes gives users the options to perform random perspective scaling, rotation, translation, flips, elastic transforms, gaussian blur, rectangular dropout, brightness/contrast adjustments, and addition of random gaussian noise (Fig.1e). The training and prediction processes take place locally using the central processing unit (CPU) and, where available, the graphics processing unit (GPU) of the host computer.

**Fig. 1:**
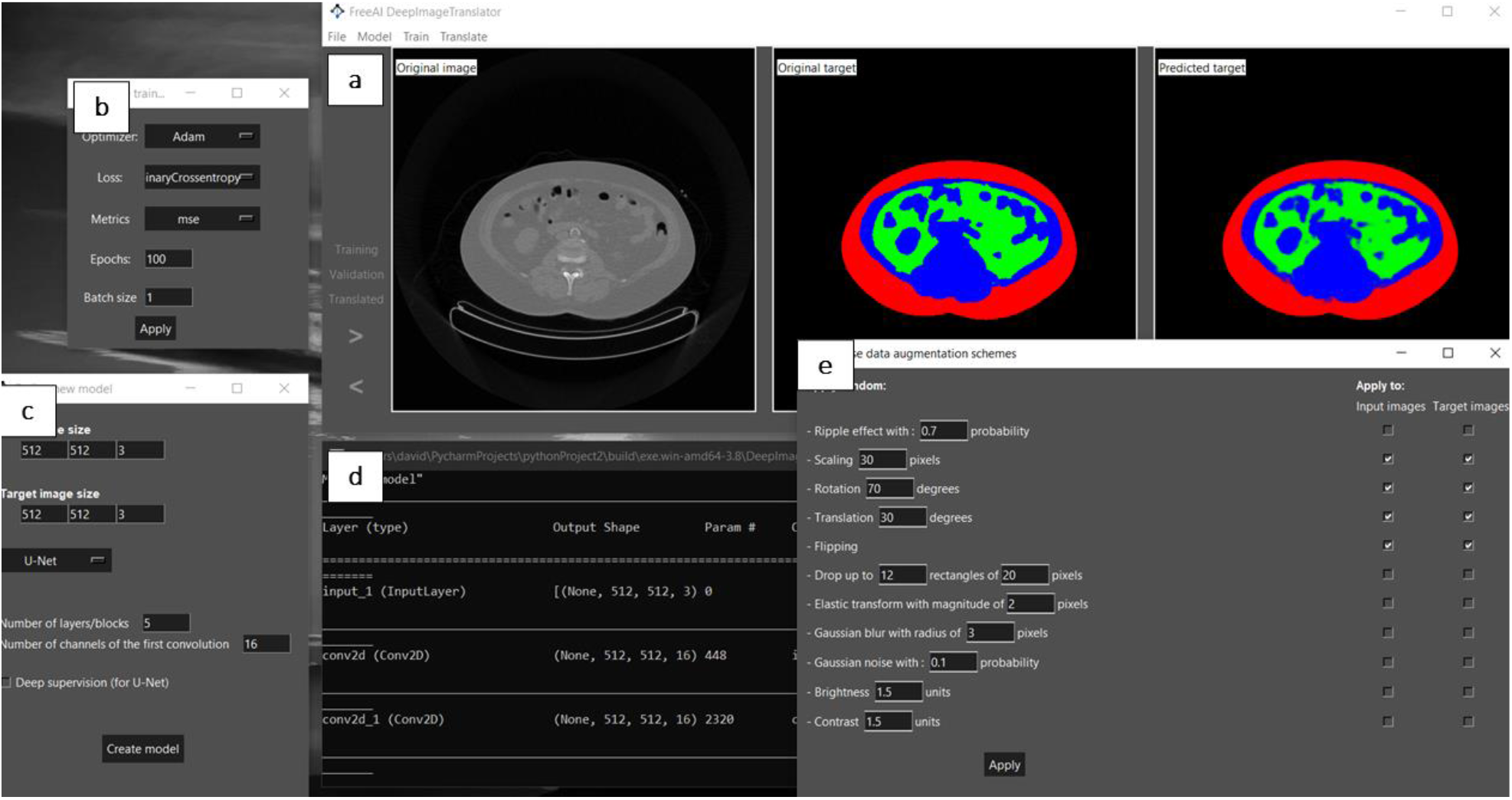
Different features of DeepImageTranslator. **a**, The main window for image viewing. **b**, Training hyperparameter selection window. **c**, Neural network model builder. **d**, Command prompt window for training monitoring. **e**, Image augmentation toolbox.

The general workflow when using DeepImageTranslator begins with the preparation of input and target images for model training, which, in the case of CT image segmentation, would consist of the original CT images and hand-labelled segmentation masks. These images are subsequently split into a training set (for model training) and a smaller validation set (to monitor the performance of the model throughout the training process). The sample size of the training set can be artificially increased through image augmentation techniques implemented in the image generator of the DeepImageTranslator. The CNN follows the general structure of U-net [18] with a few adjustments to allow for easier customization (Fig.2a). Notably, the number of layers (or depth) of the network can be specified to any value; the number of feature maps (or channels) extracted at each layer can also be changed in accordance with the complexity of the image translation task. Furthermore, instead of prescribing a predetermined and fixed image size (traditionally 128×128, 256×256, or 512×512), we implemented appropriate resizing procedures (after the 2×2 up-sampling layer and before concatenation, Fig.2a) within a CNN such that the DeepImageTranslator is compatible with input and target images of any size or dimension. Moreover, our CNN can also be deeply supervised by concatenating the output of the antepenultimate layer with the final output.

**Fig. 2:**
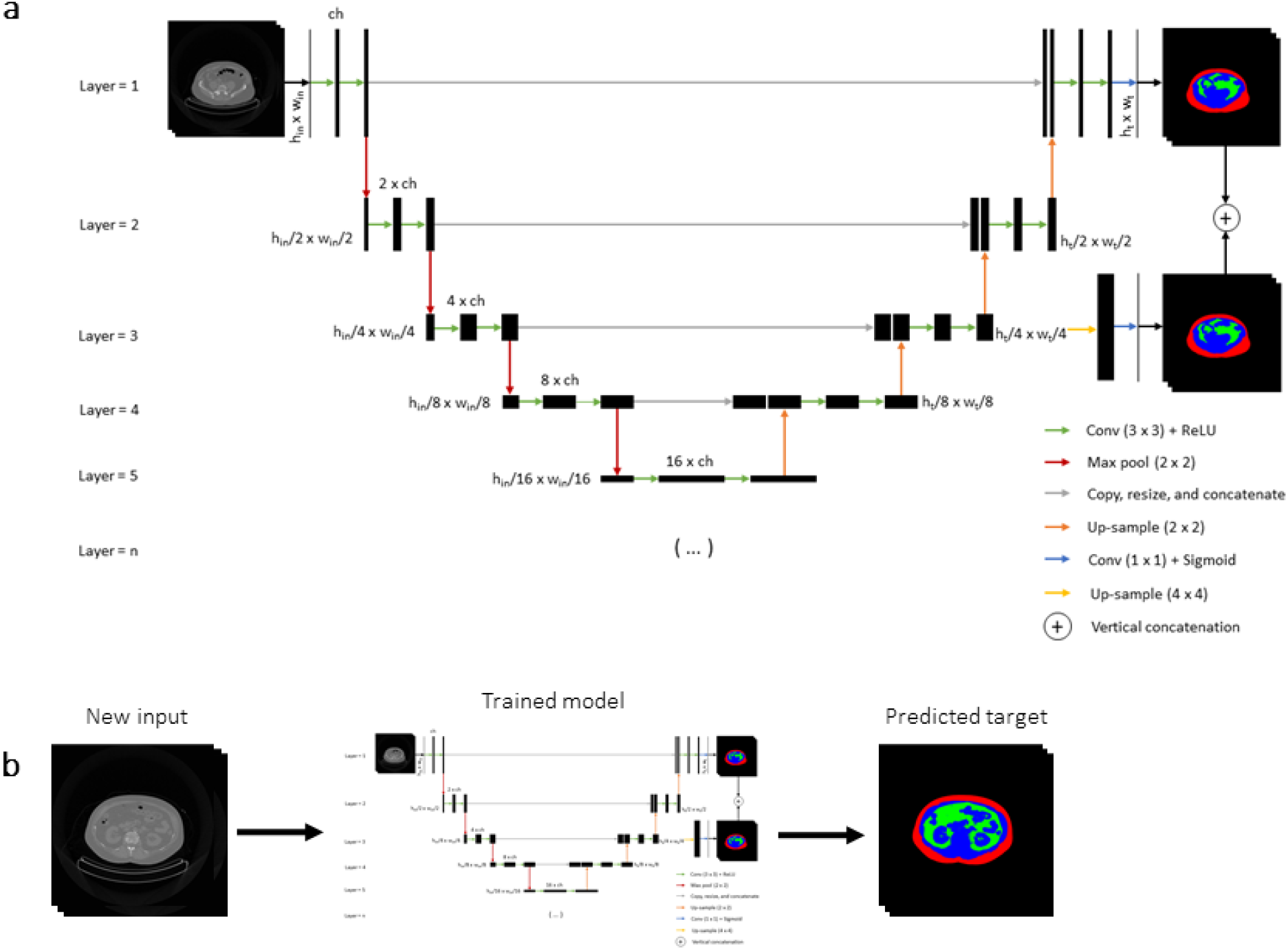
The pipeline for using the DeepImageTranslator. **a**, The construction of a custom U-net-like convolutional neural network and model training. **b**, The use of the trained neural network to make predictions based on new input data. *ch*: number of convolution maps (channels) after the first 3×3 convolution; *Conv.*: convolution; *h_in_*: input image height; *h_t_*: target image height; *ReLU*: rectified linear activation function; *w_in_*: input image width; *w_t_*: target image width.

At the end of each training epoch, model weights are automatically saved if they resulted in an improvement in model accuracy. This allows the user to reload weights of a specific epoch to either translate new images and save model predictions (Fig.2b), to continue training using the same training set, or to perform transfer learning based on a different set of training images.

### Applications of DeepImageTranslator – semantic segmentation of abdominal CT images Subject characteristics and training sample

The performance of CNNs in CT segmentation has already been demonstrated in large datasets. Therefore, in this study, we tested the accuracy and generalization ability of the DeepImageTranslator using a small dataset. Our training subjects consisted of 2 females and 1 male aged from 57 to 66 years (Table 1). A total of 165 abdominal axial images (512×512 pixels) were selected from whole-body CT scans (see Methods). These images were manually labelled using a semi-automatic approach. The bone, muscles, and other visceral organs (labelled in blue) were separated from adipose tissue, which was further segregated into the visceral (labelled in green) and the subcutaneous (labelled in red) compartments.

**Table 1:** Subject characteristics.

|  | Subject | Number of images | Sex | Weight (kg) | BMI (kg/m<sup>2</sup>) | Waist circumference (cm) |
| --- | --- | --- | --- | --- | --- | --- |
| **Three-fold cross-validation set** | 1 | 52 | F | 74.7 | 32.3 | 97.5 |
|  | 2 | 56 | F | 70.0 | 25.7 | 90.3 |
|  | 3 | 57 | M | 75.5 | 26.1 | 101.0 |
| **Testing set** | 1 | 53 | M | 110.9 | 35.0 | 122.5 |
|  | 2 | 42 | F | 58.4 | 25.3 | 81.0 |

*BMI*: body mass index; *F*: female; *M*: male.

### Model performance and effects of data augmentation with/without deep supervision

We used a three-fold cross-validation scheme to assess the accuracy of our trained models. Briefly, model training was replicated three times, each time using the images from a different subject as the validation set, leaving the images of the other two subjects in the training set. Therefore, the CNN was trained each time on data from 2 subjects, while images from the remaining subject was used for model assessment only. As expected, binary cross-entropy loss was generally lower in the training dataset than in the validation dataset and achieved a plateau by 20-40 epochs (Fig.3a and 3b).

**Fig. 3:**
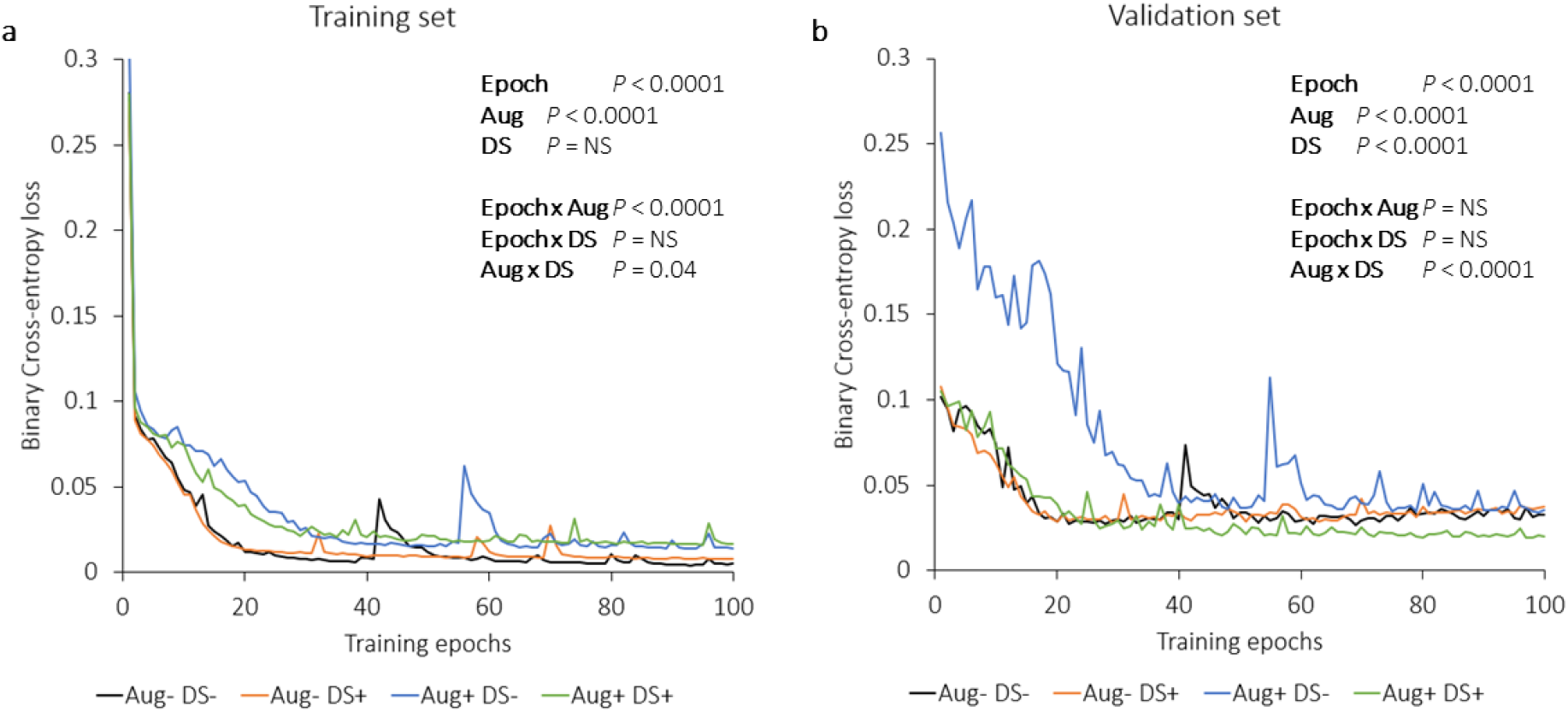
Evaluation of the effects of data augmentation with or without deep-supervision on training and validation cross-entropy loss for image segmentation. **a**, Binary cross-entropy loss across training epochs for the training split. **b**, Binary cross-entropy loss across training epochs for the validation split. *Aug*−/+: with/without data augmentation; *DS*−/+: with/without deep-supervision; *NS*: not significant. *P*-values are for 3-way ANOVA.

We first systematically evaluated the effects of data augmentation with/without deep supervision on model performance throughout the training process, both in the training set and the validation set. With data augmentation, the training cross-entropy loss was higher (three-way ANOVA *P* < 0.0001, with epoch number and deep-supervision as covariates, Fig.3a) and became closer to the validation loss. In the training dataset, deep supervision (DS) increased training speed only in the augmented (Aug) data over the first 30 epochs (Fig.3a). This differential effect of deep-supervision on training acceleration and loss reduction in augmented *vs.* non-augmented data was markedly more significant in the validation loss curves (Aug × DS interaction *P* < 0.0001). In the validation set (Fig.3b), loss curves before data augmentation were nearly superimposable whether deep-supervision was performed (Fig.3b, orange curve) or not (Fig.3b, black curve). These curves gradually increased beginning from epoch 20, again suggesting model overfitting without data augmentation after 20 epochs. However, when data augmentation was introduced, validation loss continued to decrease even after epoch 80. As expected, when augmentation was performed without simultaneous deep-supervision (Fig.3b, blue curve), training speed was significantly slowed down; as a result, by the end of 100 epochs, validation loss was similar to that without augmentation. Nonetheless, when augmentation was done concomitantly with deep supervision (Fig.3b, green curve), training speed was restored, and validation loss was then significantly lower by epoch 100.

We then used the model with the lowest validation cross-entropy loss to generate predicted targets for the validation set. The predicted segmentation map was subsequently compared with the ground truth target to calculate the Dice coefficient, model sensitivity and specificity, as well as the fractional area difference for each of the three tissue compartments (see Methods). As shown in Fig.4, data augmentation resulted in statistically significant gains in Dice coefficient, model sensitivity, and specificity. Here, color channels 1 (red), 2 (green), and 3 (blue) represent the subcutaneous adipose tissue (ScAT), visceral adipose tissue (VAT), and lean tissues, respectively. Median Dice coefficient increased by 2 to 3 percentage points in the three-fold cross-validation assessment performed with data augmentation and deep supervision compared to that without (Table 2). The best model achieved Dice coefficients of 0.99, 0.95, and 0.97 in segmenting the ScAT, VAT, and lean tissues in the validation set, which contained only images that were not used to train the model.

**Table 2:**
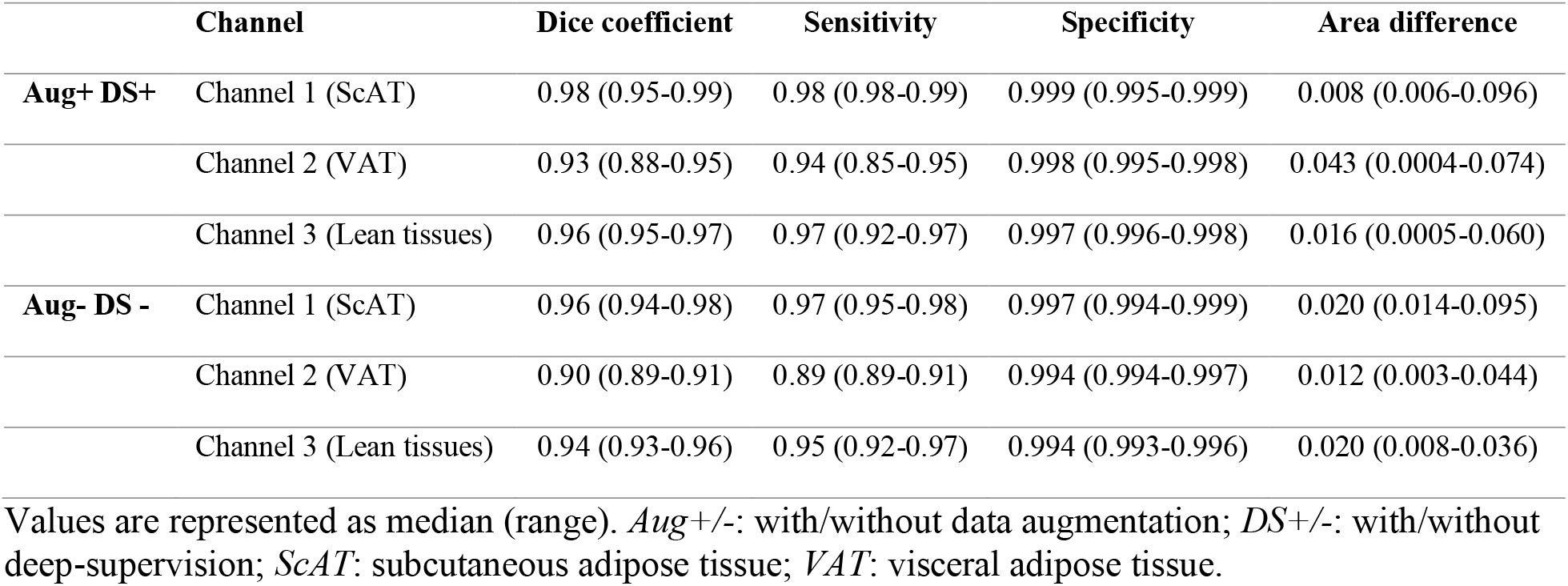
Model performance in semantic segmentation of different validation datasets in three-fold cross-validation.

**Fig. 4:**
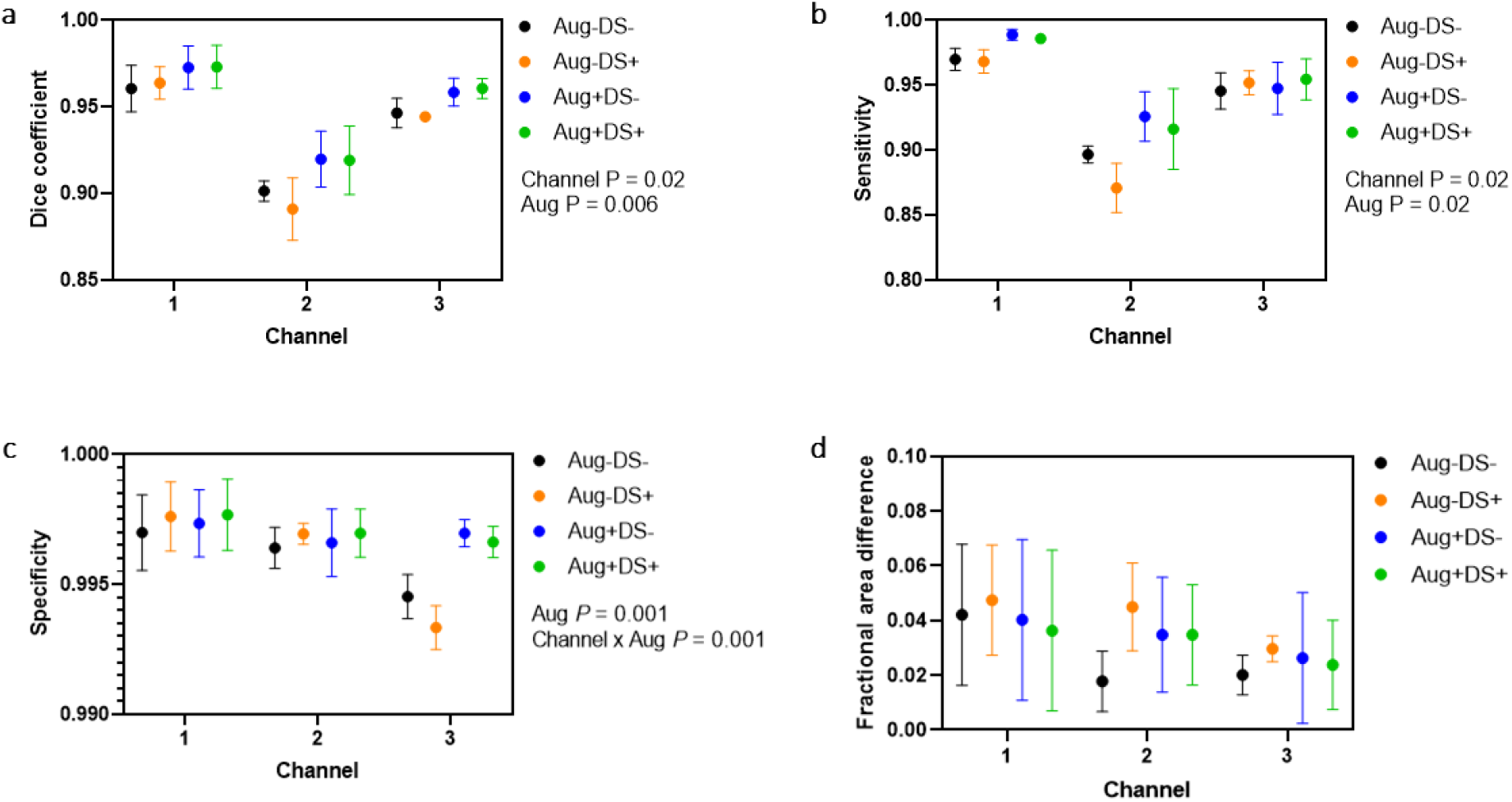
Evaluation of the effects of data augmentation with or without deep-supervision on accuracy metrics for image segmentation of the best models. **a**, Dice coefficient between ground truth maps and model predictions for each of the segmentation channels (*i.e.* tissue compartments). **b**, Sensitivity of models to detect body compartments. **c**, Specificity of models to detect tissue compartments. **d**, Fractional area difference between ground truth maps and model predictions. *Aug*−/+: with/without data augmentation; *Channel 1*: subcutaneous adipose tissue; *Channel 2*: visceral adipose tissue; *Channel 3*: other tissues; *DS*−/+: with/without deep-supervision; *NS*: not significant. *P*-values are for 3-way ANOVA.

Fig. 5a shows 3 randomly chosen CT images from the validation dataset used in each of the three cross-validation assessments, their corresponding ground truth segmentation maps, and the model predictions. White arrows point to examples of disagreement between the manually labeled map and the model prediction. Fig.5b illustrates rare occasions in which the trained neural network made obvious segmentation mistakes; many of these were due to ambiguous delineation of the abdominal wall, causing the model to label intra-abdominal compartments as extra-abdominal and *vice versa*.

**Fig. 5:**
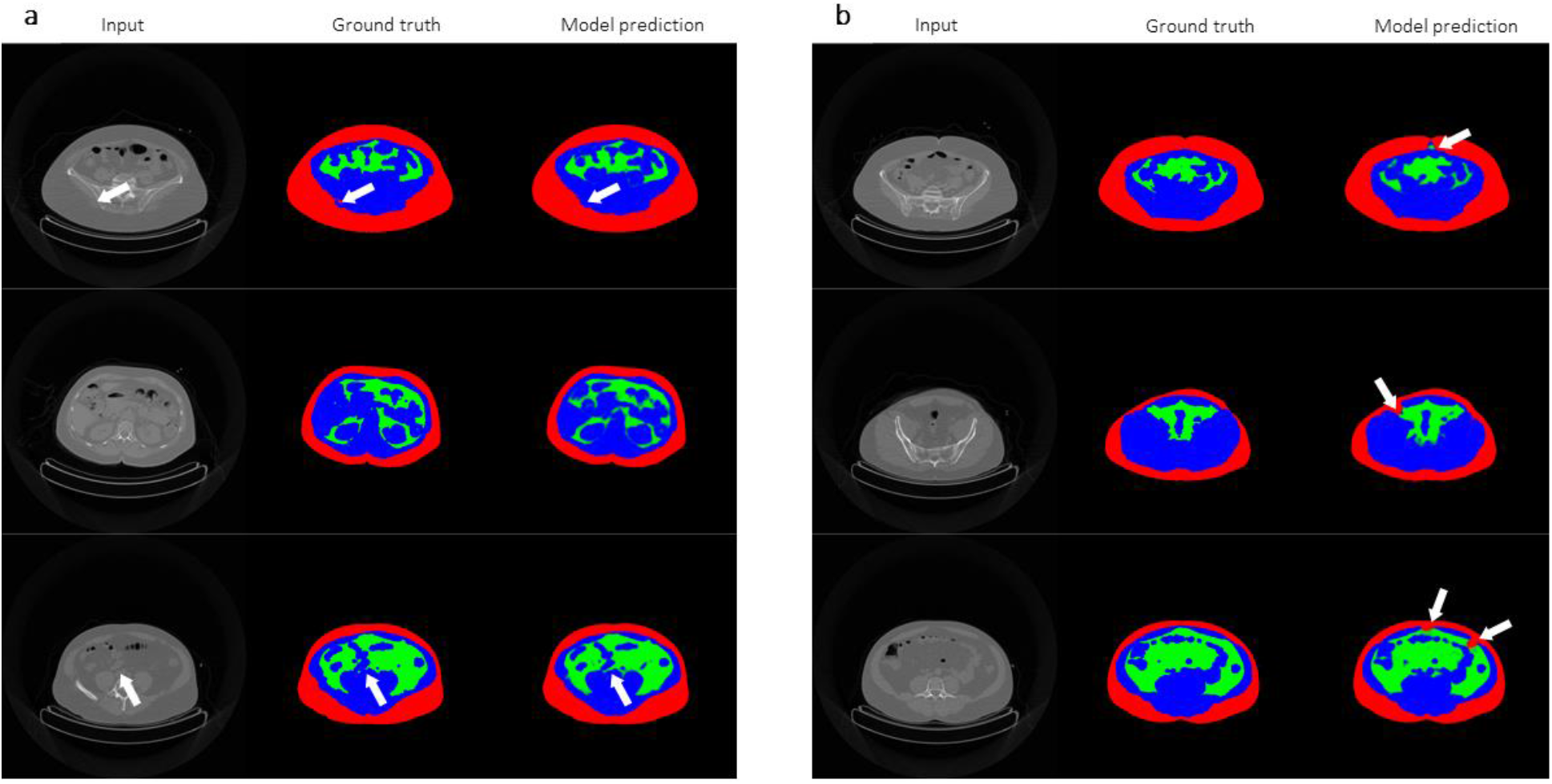
Comparison between hand-labelled segmentation maps and model predictions. **a**, *left panel*: input images randomly selected from the validation split for inclusion in the figure; *middle panel*: ground truth segmentation maps; *right panel*: model predictions based on images in the left panel. **b**, *left panel*: input images manually selected from the validation split to show rare instances of model error; *middle panel*: ground truth segmentation maps; *right panel*: model predictions based on images in the left panel. White arrows indicate area of disagreement between hand-labelled maps and model predictions.

### Generalizability and effect of sample size

Since the 165 images in our training and validation sets all came from a homogenous set of subjects that are overweight to slightly obese, we wanted to assess the generalizability of our CNN regardless of body weight and composition. Therefore, we selected 53 CT images from a male subject suffering from more severe, class II obesity, with body weight, BMI, and waist circumference of 110.9 kg, 35 kg/m^2^, and 122.5 cm, respectively. We trained a new model with the complete original training set of 165 images for 100 epochs and obtained a network capable of segmenting images from the obese subject with Dice coefficients of 0.99, 0.94, and 0.97 for ScAT, VAT, and lean tissues, respectively (Table 3 and Fig.6a).

**Table 3:**
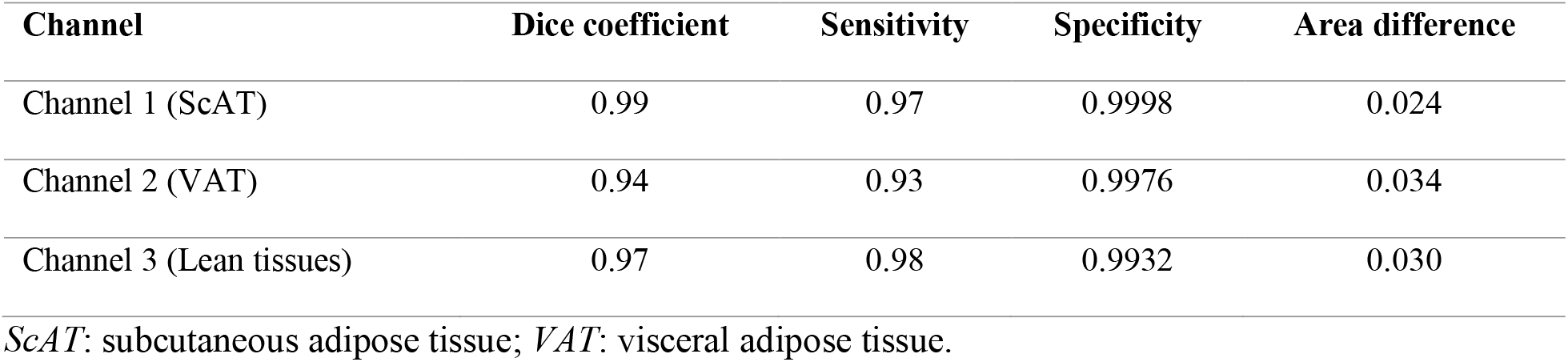
Model performance in semantic segmentation of out-of-sample CT images from an obese subject.

| Channel | Dice coefficient | Sensitivity | Specificity | Area difference |
| --- | --- | --- | --- | --- |
| Channel 1 (ScAT) | 0.99 | 0.97 | 0.9998 | 0.024 |
| Channel 2 (VAT) | 0.94 | 0.93 | 0.9976 | 0.034 |
| Channel 3 (Lean tissues) | 0.97 | 0.98 | 0.9932 | 0.030 |

ScAT: subcutaneous adipose tissue; VAT: visceral adipose tissue.

**Fig. 6:**
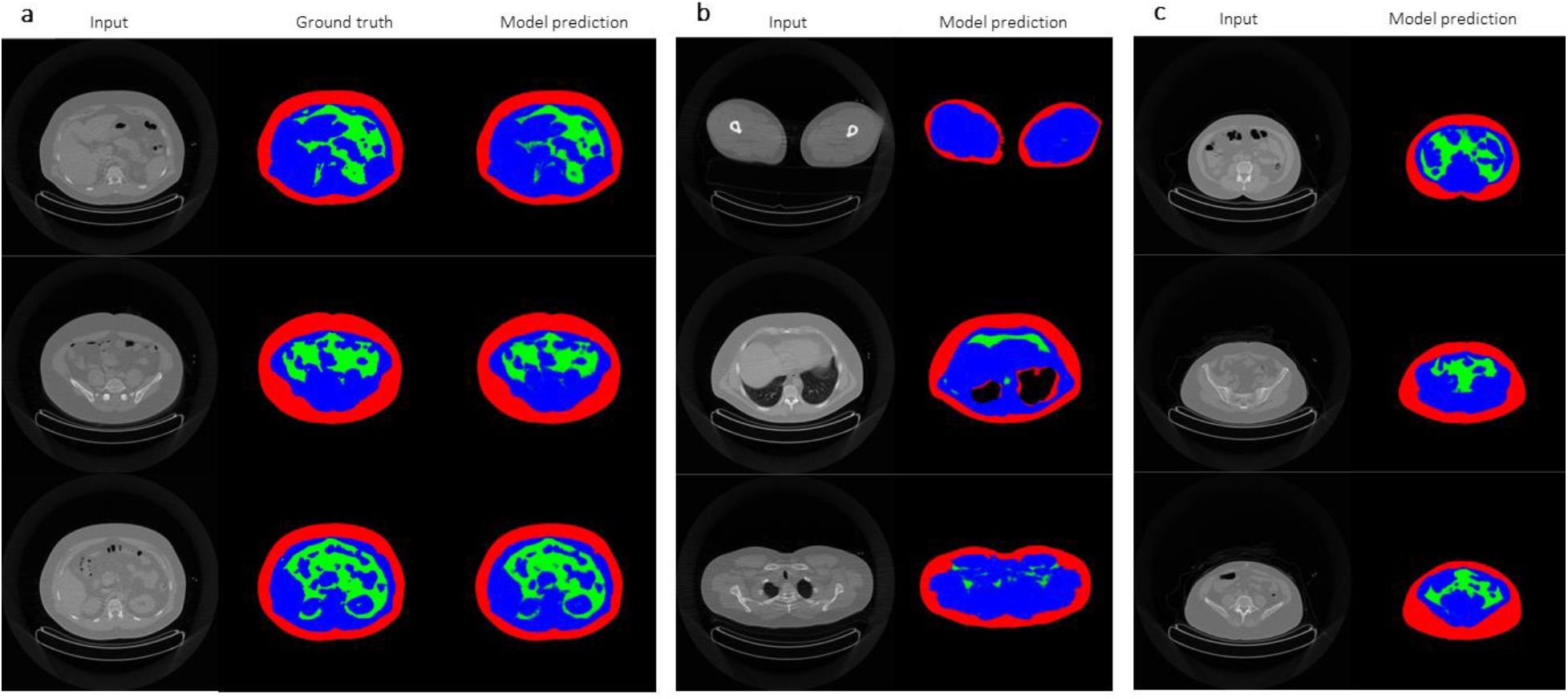
Assessment of out-of-sample model generalizability based on scans from a severely obese male subject and a very lean female subject. **a**, Model generalizability in the obese male subject. *left panel*: input images randomly selected for inclusion in the figure; *middle panel*: ground truth segmentation maps; *right panel*: model predictions based on images in the left panel. **b**, Model generalizability in the obese male subject. *first row*: image randomly chosen to from the leg area (left panel) and accompanying model segmentation (right panel); *second row*: image randomly chosen to from the thoracic area (left panel) and accompanying model segmentation (right panel), showing accurate model segmentation of the lungs; *third row*: image randomly chosen to form the shoulder area (left panel) and accompanying model segmentation (right panel). **c**, Model generalizability in the lean female subject. *left panel*: input images randomly selected for inclusion in the figure; *right panel*: model predictions based on images in the left panel.

To illustrate the generalizability of the CNN trained using DeepImageTranslator, we used our model (which was trained using only scans of the abdominal region) to segment CT images of the legs, thorax, and the upper-shoulder region of the same obese subject. Fig.6b shows randomly selected images from each of these regions and the associated model prediction. Despite never having seen images from these body areas, the model achieved remarkable generalizability by accurately labelling ScAT, and intra-thoracic adipose tissue.

Furthermore, we also tested the generalizability of our model with a very lean, female subject, with body weight, BMI, and waist circumference of 58.4 kg, 25.3 kg/m^2^, and 81.0 cm. Fig.6c shows randomly selected slices and corresponding model predictions, demonstrating excellent automatic segmentation results.

Next, we systematically determined the effect of training sample size on the generalizability of our pipeline by training distinct models with the original training set and with different fractions thereof (84 (50%), 42 (25%), 17 (10%), 10 (6%), or 5 (3%) training images). For each of these different sample sizes, we trained 3 different models, each time using a different subset of the complete training set. As shown in Fig.7a-d, the use of reduced training sets did not result in statistically significant performance loss even when sample size reached a remarkable minimum of only 17 training images (Tukey’s multiple comparisons test *P* = NS). For reference purposes, previous reports using significantly larger training datasets [3–7] showed ScAT segmentation accuracy scores that were mostly inferior to that of our model (Table 4). Furthermore, these models were trained and tested on single-slice axial abdominal CTs images centered at the L3 vertebra level only. Therefore, intraabdominal composition should be roughly similar across these training images; Dice scores for VAT segmentation were thus higher than that achieved by our volumetric models, as expected. In addition, studies using volumetric abdominal CT scans (Table 4) showed similar or inferior results for ScAT and VAT segmentation [2, 8].

**Table 4:** Previously reported Dice scores for ScAT and VAT segmentation.

| Number of training images | Dice score for ScAT segmentation | Dice score for VAT segmentation | Reference |
| --- | --- | --- | --- |
| **40 volumetric CTs (32 slices x 256 pixels x 256 pixels) with 3D U-net** | 0.95-0.96 |  | [2] |
| **2430** single-slice axial abdominal CTs centered at the L3 vertebra level | 0.98 | 0.97 | [3] |
| **3774** single-slice L3 abdominal images | 0.99 | 0.98 | [4] |
| **800** single-slice images at the L3/L4 level | 0.98 | 0.96 | [5] |
| **180** axial CT images at the supra-acetabular level | 0.97 |  | [6] |
| **883** images at the inferior endplate level of the 3rd lumbar vertebra | 0.97 | 0.97 | [7] |
| **120** volumetric CT images | 0.98 | 0.92 | [8] |

ScAT: subcutaneous adipose tissue; VAT: visceral adipose tissue

**Fig. 7:**
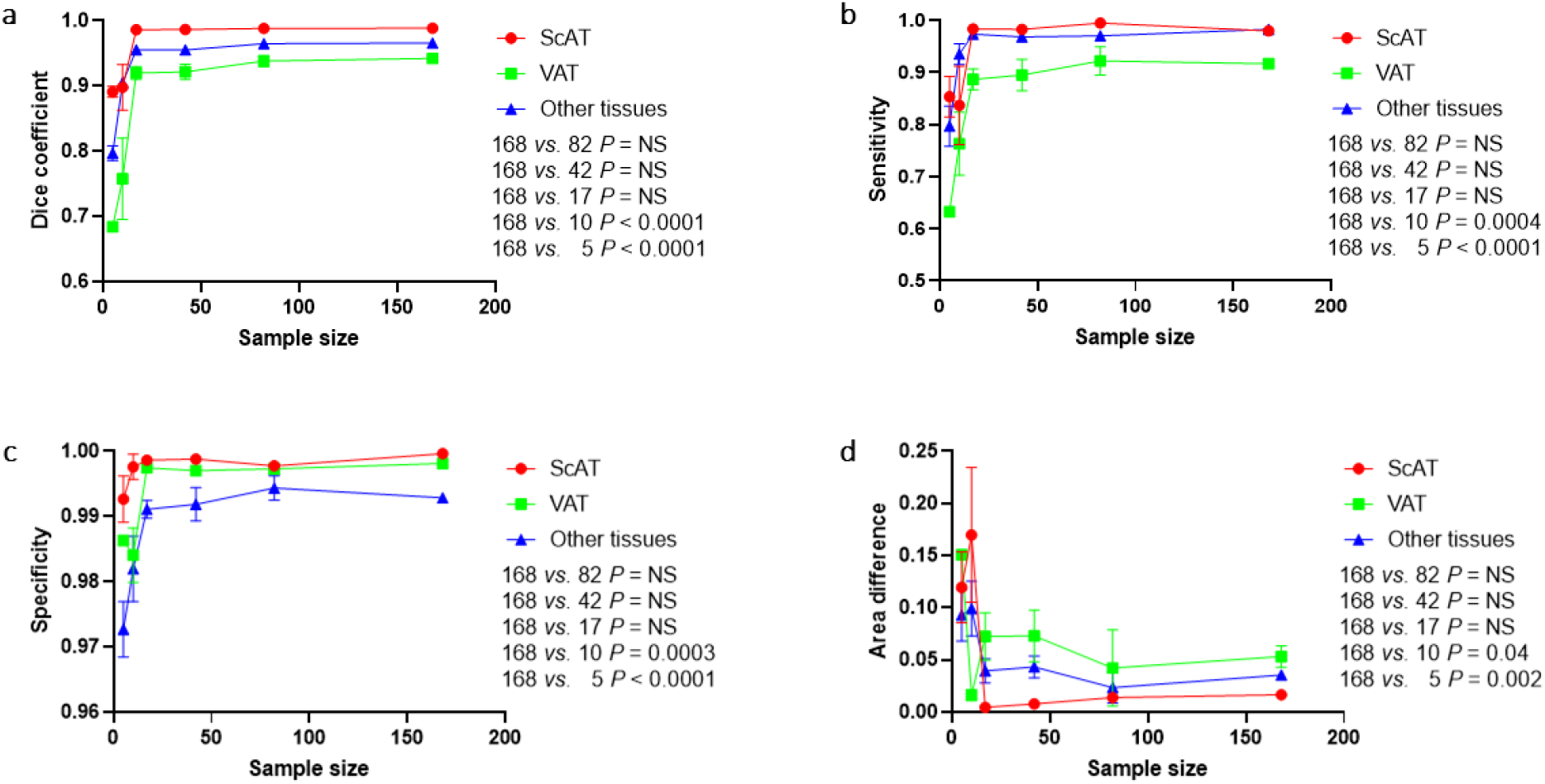
Evaluation of sample size. **a**, Dice coefficient between ground truth maps and model predictions for each of the segmentation channels (*i.e.* tissue compartments). **b**, Sensitivity of models to detect tissue compartments. **c**, Specificity of models to detect tissue compartments. **d**, Fractional area difference between ground truth maps and model predictions. *ScAT*: subcutaneous adipose tissue (red circle 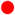); *VAT*: visceral adipose tissue (green square 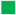); Other tissues (blue triangle 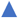). *P*-values between different sample sizes are for Tukey’s multiple comparisons tests.

### Applications of DeepImageTranslator – noise reduction for thoracic CT images

To further demonstrate the versatility of DeepImageTranslator in entirely different image translation tasks, we evaluated the software’s performance in thoracic CT image noise reduction. During CT examinations, noise can be introduced at various stages of acquisition and transmission. Here, we created noisy CT images using additive Gaussian noise model [9, 19, 20]. To train neural networks for noise removal, we first extracted CT slices from the thoracic area of our 3 original subjects. We proceeded with the same three-fold cross validation scheme as described above in the image segmentation section.

Our models achieved excellent noise-removal results even with only 100 epochs of training using the mean average error (MAE) as the loss function (Fig.8). However, since the validation curves for MAE, mean squared error (MSE), structural similarity index measure (SSIM), and peak signal-to-noise ratio (PSNR) were still improving at epoch 100, we randomly selected one of the three models and continued training for 200 epochs. This allowed our final model to reach MAE, MSE, SSIM, and PSNR values of 0.002, 0.000004, 0.993, and 44.8 on validation images.

**Fig. 8:**
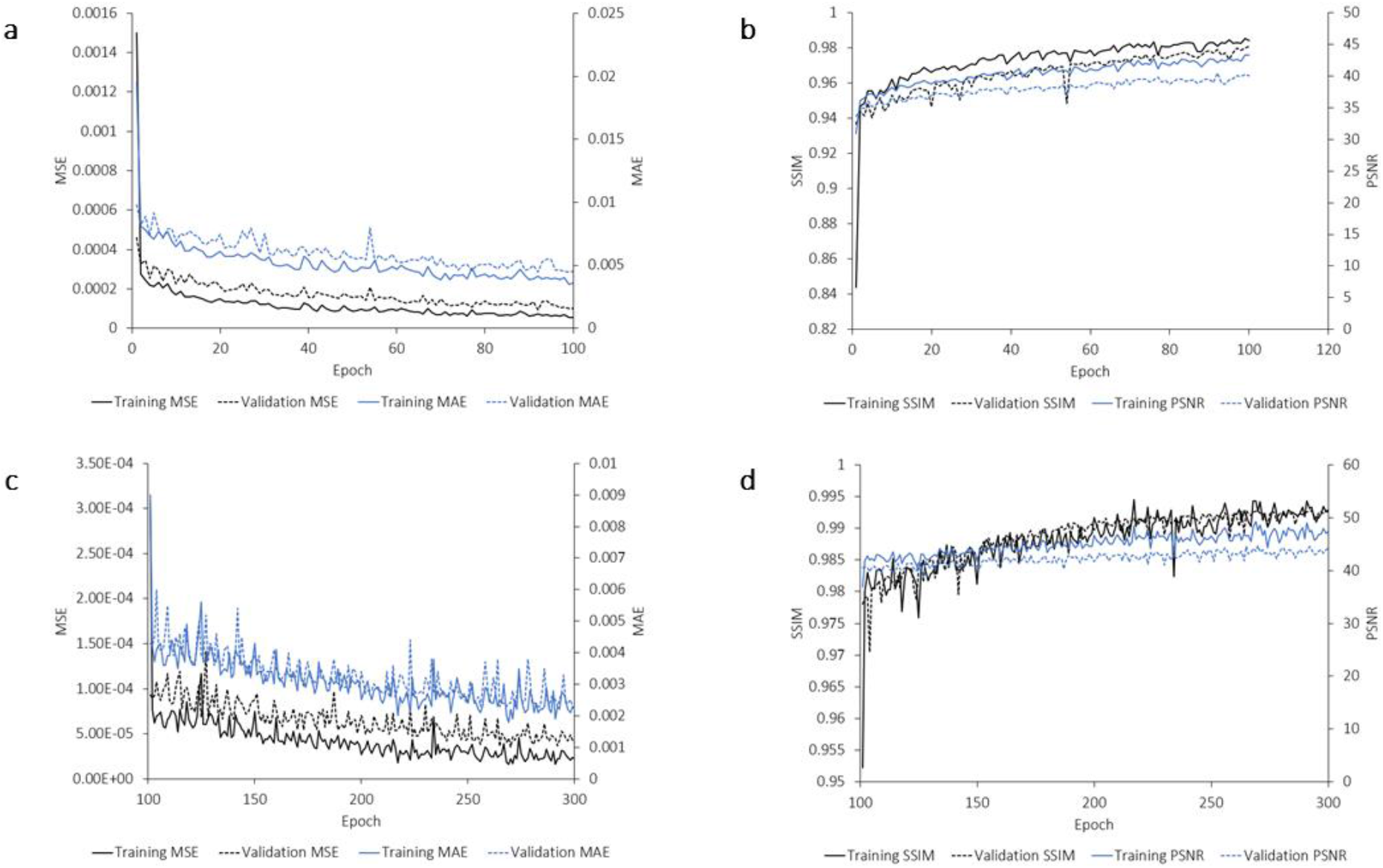
Model efficiency for noise reduction. **a**, Evolution of training (solid lines) and validation (dashed lines) mean square error (MSE) and mean average error (MAE) across the first 100 epochs with three-fold cross-validation. **b,** Evolution of training and validation structural similarity index measure (SSIM) and peak signal-to-noise ratio (PSNR) across the first 100 epochs with three-fold cross-validation. **c,** Evolution of training and validation MSE and MAE across the last 200 epochs with one randomly selected model. **d,** Evolution of training and validation SSIM and PSNR across the last 200 epochs with one randomly selected model. *MAE*: mean average error; *MSE*: mean square error; *PSNR*: peak signal-to-noise ratio; *SSIM*: structural similarity index measure.

As shown in Fig.9a, our model was able to considerably reduce the noise introduced in the validation images (left panels) to produce predictions (right panels) that were almost indistinguishable from the original (noiseless) images (middle panels). Fig.9b shows enlargements at the right and left lungs and demonstrates our CNN’s ability to accurately recover detailed pulmonary vasculature invisible on the noisy images.

**Fig. 9:**
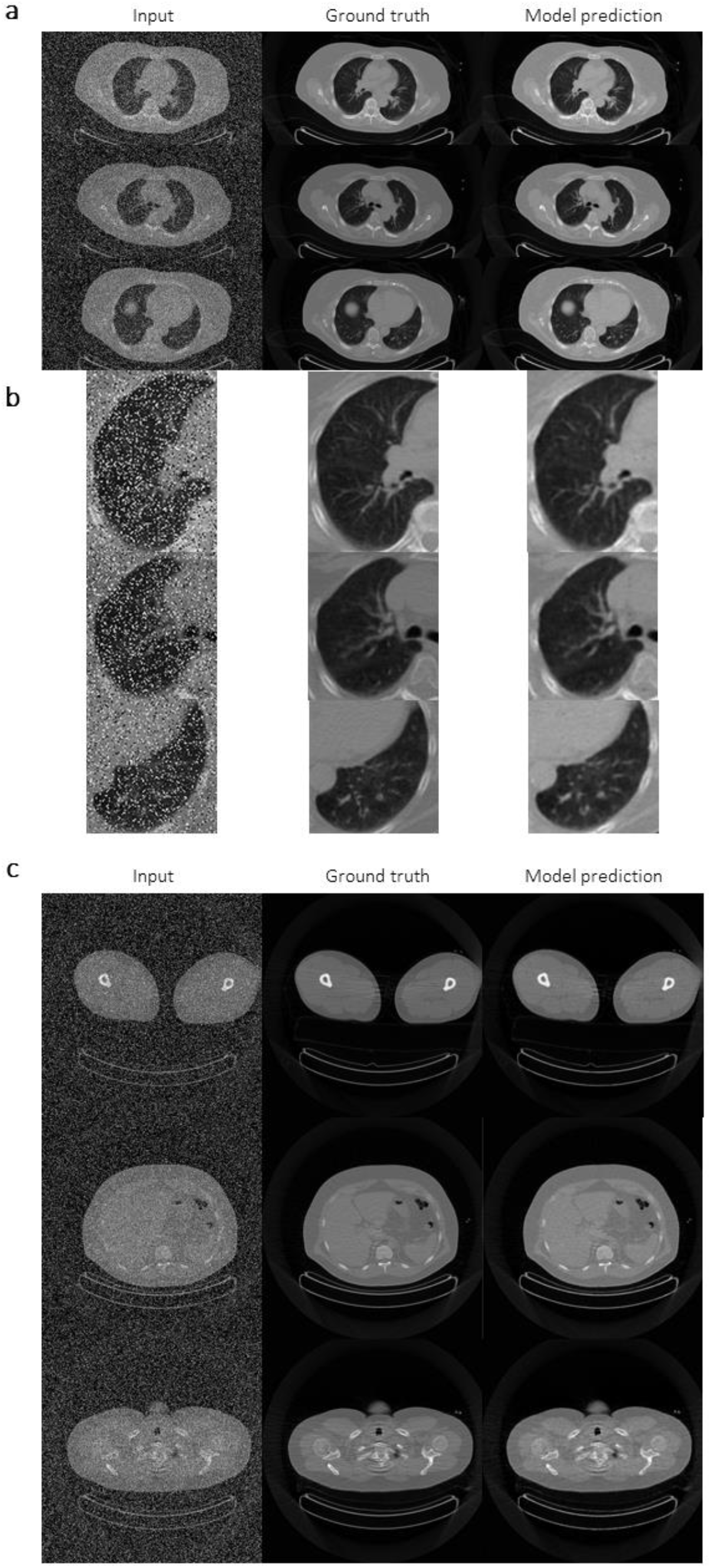
Assessment of model performance and generalizability for noise reduction. **a**, Randomly chosen slides from the validation split showing model performance in noise reduction comparing the noisy input images (left panel), original (noiseless) images, and image generated by the model (right panel) based on the noisy input images. **b**, Details of the right and left lungs of the images shown in panel **a**, demonstrating our model’s ability to recover fine details of the pulmonary vasculature. **c**, Randomly selected slices at the level of the legs, abdomen, and shoulder from the out-of-sample obese subject, showing generalizability of our trained model. *Left panel*: noisy CT images; *middle panel*: original (noiseless) images; *right panel*: model prediction based on images from the left panel.

### Generalizability

To establish the generalizability of our neural network, we used the model to denoise whole-body scans from our obese subject. Fig.9c shows randomly selected slices from the leg, abdominal, and the shoulder areas and illustrate our network’s ability to generalize in different regions of the body despite being trained only on thoracic scans. Furthermore, we also increased the noise level by approximately 60%, using thoracic scans of the same obese subject, to the point of rendering the CT images completely unreadable; we then demonstrated that our trained model was capable of removing all the noise (Fig.10a) and of recovering fine pulmonary vascular markings (Fig.10b).

**Fig. 10:**
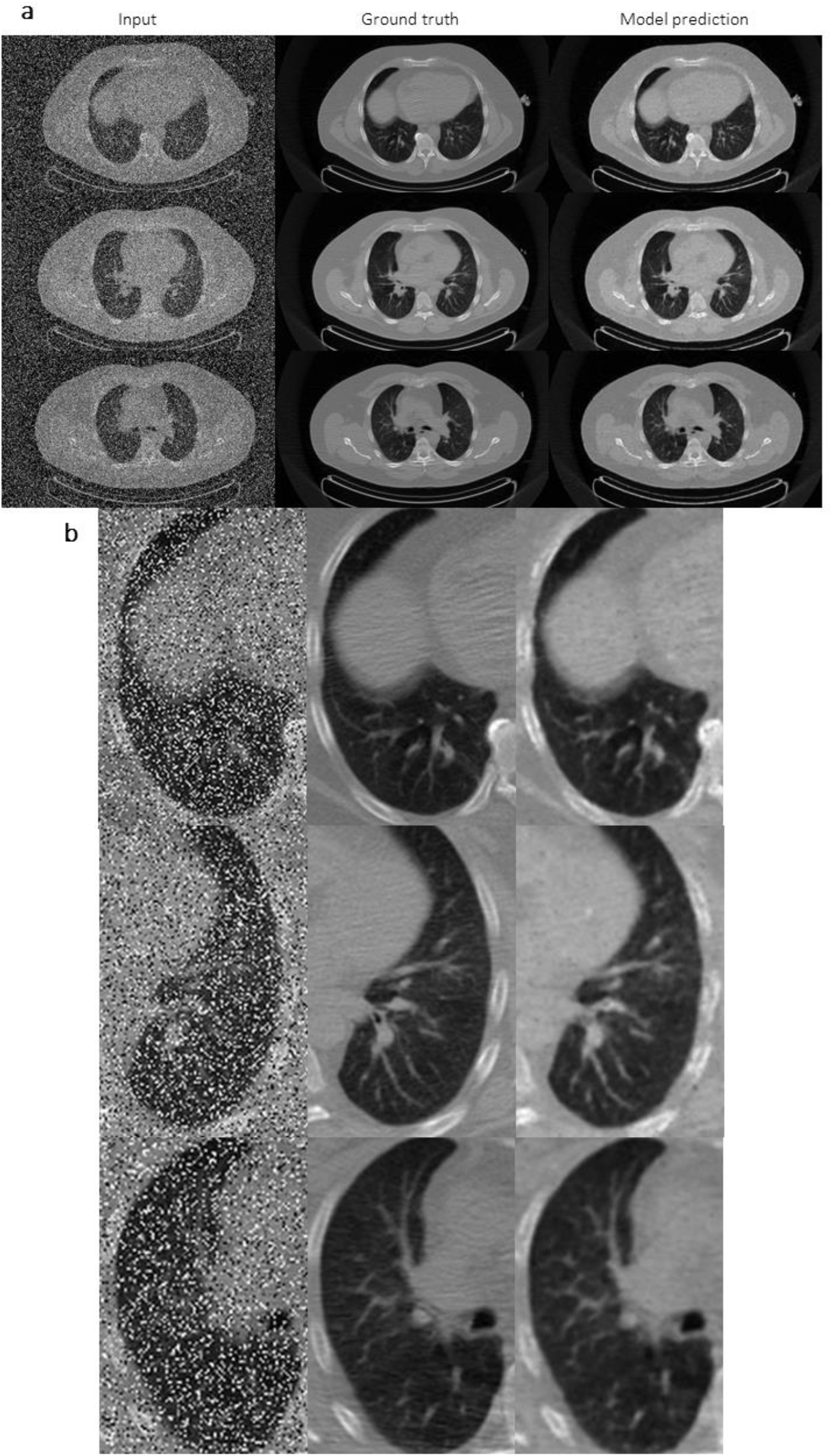
Assessment of model generalizability for increased noise level. **a**, Randomly chosen thoracic CT slices from the obese subject with an increase of approximately 60% in noise intensity, demonstrating the efficiency of our model for noise reduction. **b**, Enlargement at the right and left lungs of the images shown in panel **a**, demonstrating our model’s ability to recover fine details of the pulmonary vasculature. *Left panel*: noisy CT images; *middle panel*: original (noiseless) images; *right panel*: model prediction based on images from the left panel.

### Training duration and hardware requirements

Finally, to test training duration using different types of hardware, we calculated the time to complete one epoch of 18 CT segmentation training images and 9 validation images. Training time for one epoch was approximately 5 minutes on a Windows laptop with 8 Gb of random-access memory (RAM) (AMD A4-9125 RADEON R3 processor) and 90 seconds on a MacBook Air with 8 Gb of RAM (Intel core i5 processor) from early 2015. This would allow the model to achieve the highly accurate segmentation results shown in Fig.7 in 17 hours and 5 hours with these two devices, respectively. Using an NVIDIA GeForce RTX 2060 GPU for massively parallel computing, training time was reduced to 3 second/epoch; 200 epochs can thus be completed under 11 minutes.

## DISCUSSION

DeepImageTranslator is designed to be a user-friendly graphical interface tool that allows researchers with no programming experience to easily build, train, and evaluate CNNs for image translation. Compared to existing software programs, our tool also allows users to customize their CNN (*e.g.* number of layers/channels, use of deep-supervision, and input layer resolution), the type of model optimizer algorithm, the loss function, and the data augmentation schemes. We showed that using only a standard personal computer, it is possible to train neural networks for accurate semantic segmentation of CT images. This free, open-source tool is available at: (will be uploaded with the publication of this article) for Windows 10. It is actively maintained by the first author of this work and is distributed under the GNU General Public License (version 3.0). Future directions for our tool include the addition of new data augmentation methods and of different CNN architectures such as cycle consistent generative adversarial networks (cGAN), which would allow translation using unpaired images and targets.

In 3D semantic segmentation of CT images of the abdominal area, our trained models achieved accuracy levels comparable or superior to those reported elsewhere using different neural networks [2–8] and using traditional methods [21]. Our deep-learning tool analyses CT examinations on a slide-by-slide basis. Of course, it is possible to extend 2D inputs into 3D inputs whereby the model receives the entire 3D CT dataset or multiple slices from a given patient. Nevertheless, we did not implement such an approach since our models have already achieved high accuracy without this type of extension. Furthermore, using three-dimensional inputs will multiply the computational demands and curtail the flexibility of our deep-learning pipeline.

We performed three-fold cross-validation procedures to systematically assess the effects of data augmentation with or without deep-supervision. Our results showed that across the first 100 training epochs, data augmentation resulted in statistically significant reductions in validation loss only when accompanied by deep-supervision, which curbed the effect of data augmentation on training speed. Therefore, we would recommend the implementation of image augmentation schemes with concomitant increase in training duration and/or techniques to accelerate training. Previous reports on semantic CT segmentation used relatively large training datasets. Therefore, we aimed to determine the minimum sample size required to achieve high predictive accuracy and model generalizability in a small dataset using the DeepImageTranslator. We found that, with appropriate data augmentation, 17 image-target pairs were sufficient to obtain high segmentation accuracy in test images. Furthermore, our segmentation model was also able to generalize to body areas that they were not trained on and independently of body weight and composition. Therefore, for studies with small datasets, researchers can randomly select a very small subset of images for manual labeling, which can then be used to train a specialized CNN model with DeepImageTranslator to fully automate segmentation of the entire dataset, thereby saving tremendous time and labor.

In the CT image noise reduction task, models trained with DeepImageTranslator also showed comparable to superior performance compared to previously reported deep-learning [9–11] and non-deep-learning methods [19, 20]. However, it should be noted that direct comparisons of MSE, SSIM, and PSNR scores across studies are difficult due to wide methodological differences in noise simulation techniques. To simulate very noisy CT scans, we reduced image quality to very low levels and showed that our model recovered detailed vasculature of the lungs and was able to denoise whole-body scans despite having been trained only on thoracic images.

## ACKNOWLEDGMENTS

We would like to acknowledge the contribution of Éric Lavallée, Maude Gérard, Caroll-Lynn Thibodeau and Diane Lessard for their technical support and the contribution of En Zhou Ye who developed the DeepImageTranslator software. We would also like to thank Pr. Pierre-Marc Jodoin for his advice concerning the structure and methodology of this paper.

## FUNDING

This work was supported by a grant from the Canadian Institutes of Health Research (CIHR operating grants MOP53094, grant number 341582). ACC holds the Canada Research Chair in Molecular Imaging of Diabetes.

## DISCLOSURES

The authors declare no competing interests

ACC received funding by Eli Lilly, HLS Therapeutics, Janssen Inc., Novartis Pharmaceuticals Canada Inc., and Novo Nordisk Canada Inc. as a consultant. None of these commercial relationships are relevant to the present study.

## Notes

https://sourceforge.net/projects/deepimagetranslator/

https://github.com/runzhouye/DeepImageTranslator

https://figshare.com/s/b9caed5d78f9debdcd21

